# Refined cell transfer model reveals roles for *Ascl2* and *Cxcr3* in splenic localization of mouse NK cells during virus infection

**DOI:** 10.1101/2025.04.05.647391

**Authors:** Laura M. Canaday, Andrew Cox, H. Alex Feldman, Harsha Seelamneni, Ayad Ali, Jasmine A. Tuazon, Lorena Botero Calderon, Sierra N. Bennett, Allison Yan, Megan Wilson, Vijayakumar Velu, Stephen N. Waggoner

## Abstract

Cell transfer experiments complement the rigorous investigation of antiviral and antitumor functions of natural killer (NK) cells. Success in these endeavors is enhanced by expansion of small numbers of input NK cells driven by viral antigens or homeostatic proliferation in immunodeficient hosts. In contrast, analysis of other NK-cell functions, including immunoregulation, are non-proliferative and require an intact immune system in recipient mice. We reveal poor persistence of conventional congenic (CD45.1) BoyJ NK cells following adoptive transfer in comparison to CRISPR-generated CD45.1+ (JAXBoy) NK cells. Reciprocal transfers between C57BL/6 and JAXBoy mice substantially improve seeding and maintenance of donor NK cells. Using this system, we confirm that CXCR3 re-positions NK cells in the white pulp of the spleen after infection, which is vital for immunoregulation. Moreover, we discovered that the transcription factor ASCL2 is required for recruitment of NK cells into the spleen and white pulp. These results provide improved tools and novel insights into NK cell biology.

**Key points:**

- JAXBoy are superior to conventional BoyJ mice for NK cell persistence after transfer.
- CXCR3 repositions donor NK cells in T/B-zones of the spleen after infection.
- The transcription factor ASCL2 is required for NK-cell recruitment to spleen white pulp.

## Introduction

Natural killer (NK) cells are innate lymphocytes with critical roles in the killing of virus-infected cells and cancers [1, 2]. NK cells also determine the magnitude and quality of T and B cell responses via perforin-dependent killing of a subset of activated CD4+ T cells [3-5]. Enrichment of NK cells within the T- and B-cell rich white pulp regions of lymphoid tissues after infection is essential for this immunoregulation [6, 7]. Specifically, infection results in elevated expression of CXCR3 ligands in the white pulp that promotes transient re-localization of NK cells into T-cell zones, thereby facilitating physical interactions between NK cells and target T cells.

Investigations of NK cell immunoregulatory activity have largely depended upon genetic or antibody-mediated depletion of NK cells or on the generation of mixed bone marrow chimeras [8-15]. These methods aid in demonstrating the *necessity* of NK cells or specific NK-cell mediators in immunoregulation but are inadequate for ‘rescue’ studies that show *sufficiency* of these cells or genes of interest. Adoptive transfer of NK cells between mice would be an ideal tool to evaluate rescue of immunoregulatory activities. However, traditional adoptive transfer systems are complicated by high cell count requirements and limited proliferation or survival of donor NK cells in recipient animals with intact immune systems.

Here, we developed a protocol for the adoptive transfer of NK cells into immunocompetent animals that is based on CRISPR-generated congenic mice (JAXBoy) that overcome genetic issues with conventional congenic BoyJ mice models [16]. We use this system to define the necessity of the chemokine receptor CXCR3 and the migration-associated transcription factor ASCL2 in the enrichment of NK cells in the splenic white pulp after infection. Our results also reinforce the inaccuracy of presumptions regarding the degree of genetic differences between C57BL/6 and congenic BoyJ mice commonly used in cell transplantation studies [16, 17].

## Methods

### Mice

C57BL/6 (Strain #000664), B6.SJL-*Ptprc*^*a*^*Pepc*^*b*^/BoyJ (BoyJ) (Strain #002014), C57BL/6J-*Ptprc*^*em6Lutzy*^/J (JAXBoy) (Strain #033076), and B6.129P2-*Cxcr3*^*tm1Dgen*^/J (CXCR3^KO^) (Strain #005796) mice were purchased from Jackson Laboratory (Bar Harbor, ME). *Ncr1-iCre* mice were a gift from Eric Vivier (Marseille, France) and *Ascl2-floxed* mice (See details below) were a gift from Hans Clevers (Utrecht, Netherlands) and James Bridges [18, 19]. *Ncr1-iCre* mice were bred with *Ascl2-floxed* mice and littermates were used in experiments. Both male and female mice between 8-20 weeks of age were routinely utilized in experiments with no apparent sexual dimorphism in resulting data, thus both sexes were combined in data presentation. Mice were housed under barrier conditions and experiments performed under ethical guidelines approved by the Institutional Animal Care and Use Committees of Cincinnati Children’s Hospital Medical Center in compliance with the U.S. Department of Health and Human Services Guide for the Care and Use of Laboratory Animals. Staff responsible for processing samples and acquiring data were blinded to the genotype and treatment status of experimental groups of animals.

### Ascl2-floxed mice

Genetic editing was initially performed in 129/Ola-derived IB10 embryonic stem cells before extensive in-house backcrossing of *Ascl2-floxed* mice (8 generations) onto the C57BL/6 genetic background. Key NK cell receptor expression was determined by flow cytometry analysis of blood or spleen cells using the following Biolegend antibodies: Ly49H (3D10)- PE/Cyanine7, NK-1.1 (PK136)- BV421, and CD3ε (500A2)- FITC. Taconic Biosciences SNP Genotyping service was used on DNA from a female *Ascl2*-floxed mouse to assess degree of fixation to the C57BL/6J genetic background.

### Ascl2 and Cre genotyping

Genomic DNA was extracted from tail clips using the HotSHOT DNA extraction protocol (Bento Bioworks). PCR was performed with the following primers: *Ascl2* flox forward [GCG AAG CAG TAA GGG AGA CAC G], *Ascl2* flox revere [AGA ACC TGC CCG CCG TGA C], *iCre* forward [TTC TCT GAA CAC ACC TGG AAG ATG], and *iCre* reverse [AGA GTT CTC CAT CAG GGA TCT GAC]. For *Ascl2* PCR, reactions combined 2 µL DNA, 1 µL of each 10 µM primer, and 10 µL of 2X Econotaq Plus Green mastermix (Biosearch Technologies, Cat#F93481-1) in a total volume of 20 µL. The PCR cycling conditions were 94°C for 3 minutes, followed by 30 cycles of 94°C for 30 seconds, 60°C for 30 seconds, and 72°C for 1 minute, with a final extension at 72°C for 7 minutes. PCR products were separated on a 2% agarose gel and visualized under UV light, with the floxed allele at 433 bp and the wild-type allele at 383 bp. For *Ncr1-iCre* PCR, Reactions combined 1 µL DNA, 2 µL of each 10 µM primer, and 6 µL of 2X Econotaq Plus Green mastermix (Biosearch Technologies, Cat#F93481-1) in a total volume of 20 µL. The PCR cycling conditions were 95°C for 90 seconds, followed by 32 cycles of 95°C for 30 seconds, 63°C for 20 seconds, and 72°C for 30 seconds, with a final extension at 72°C for 30 seconds. PCR products were separated on a 2% agarose gel and visualized under UV light, with the KI at 348 bp.

### Confirmation of selective Ascl2 targeting in NK cells

NK cells were enriched from *Cre*(+) control, *Cre*(+) *Ascl2*^*fl/WT*^ and *Cre*(-) *Ascl2*^*fl/WT*^ littermate mice using the anti-NKp46 MicroBead Kit (Miltenyi, Cat#130-095-390) according to the kit protocol. Non-NK cells were obtained from the column flowthrough. A subset of cells was taken for pre- and post-sorting purity checks using the following Biolegend antibodies: CD335 (29A1.4)- BV605, CD3ε (500A2)- FITC, and Zombie NIR Fixable Viability Kit. Remaining enriched NK cells were then immediately lysed to extract RNA using the RNeasy mini kit (Qiagen, Cat#74104). RNA was reverse transcribed into cDNA using the iScript Reverse Transcription Supermix for RT-qPCR kit (BioRad, Cat#1708840). RT-qPCR was performed using TaqMan Gene Expression Assays to detect *Ascl2* (Mm01962673_s1) and *Gapdh* (Mm99999915_g1). Reactions were run on QuantStudio 3 and 5 Real-Time PCR Systems (ThermoFisher). *Ascl2* Ct was normalized to *Gapdh* Ct to determine ΔCt. Then the NK cell ΔCt was normalized to non-NK ΔCt for each mouse, yielding ΔΔCt. The fold-change in transcripts was then quantified (2^-ΔΔCt^).

### Adoptive transfer of NK cells

JAXBoy, BoyJ, or C57BL/6 spleens were harvested and processed into single cell suspensions (see tissue processing) from which NK cells were column enriched by depletion of non-NK cells per manufacturer’s instructions (Miltenyi Biotec, Germany) to >90% purity. One million NK cells in 100 μL Hank’s Balanced Saline Solution (HBSS) were injected retro-orbitally with a 28G needle (BD Biosciences, San Jose, CA) into C57BL/6 or JAXBoy recipient mice. Transfers were done from donor mice of same sex as recipient mice.

### Virus injection

Mice were infected with the Armstrong strain of LCMV via intraperitoneal injection of 5×10^4^ plaque forming units per mouse in 200 μL of HBSS using a 28G needle. Viral stocks were previously generated in house via propagation on BHK21 cells with viral titers determined using Vero cells [20].

### Tissue processing and cell staining for flow cytometry

Spleens, lungs, and livers were harvested from mice following euthanasia. Single-cell leukocyte suspensions were prepared from spleens by mechanical homogenization of tissues between frosted glass microscope slides (VWR, Radnor, PA) and filtration through a 70 μm nylon mesh strainer with RPMI medium 1640 + 10% fetal bovine serum + 50,000U penicillin-streptomycin (Gibco, Waltham, MA) + 2mM L-glutamine (Gibco, Waltham, MA). Lungs were homogenized by crushing with the rubber end of a 10 mL syringe through a 70 μm nylon mesh with RPMI medium 1640 + 10% fetal bovine serum + 50,000U penicillin-streptomycin + 2mM L-glutamine. Livers were homogenized with surgical scissors in RPMI + 10% fetal bovine serum. After resuspension, cells were centrifuged at 300 xg for 10 minutes. Cell pellets were homogenized in a 33% Percoll solution diluted in RPMI medium 1640. All cell suspensions were then subjected to red blood cell lysis via ACK lysing buffer for 5 minutes at 37°C. Following lysis of red blood cells, cells were plated at 2×10^6^ per well in 96-well round-bottom plates and immediately subjected to flow staining. Leukocytes were washed and resuspended in FACS buffer (Hanks Buffered Salt Solution + 5% fetal bovine serum + 0.5 mM EDTA), stained for 5 minutes at room temp with a live/dead stain, washed twice with FACS buffer, and nonspecific Fc receptor-mediated binding was then blocked via a five-minute incubation at 4°C in anti-mouse CD16/CD32 (BD BioSciences, San Jose, CA). Cells were stained with the following antibodies: CD4 (GK1.5)- BV737 [1:200], CD8α (53-6.7)- PE-CF594 [1:200], CD19 (1D3-CD19)-PerCP-Cy5.5 [1:200], CD49a (HMa1)- APC [1:100], NK1.1 (PK136)- BV711 [1:200], CD49b (DX5)- PE [1:100], NKp46 (29A1.4)- BV605 [1:50], CD45.1 (A20)- AF700 [1:100], CD45.2 (104)- AF488 [1:100], CD11b (M1/70)- BV786 [1:100], CD27 (LG.3A10)- BV421 [1:100], and live/dead-Zombie UV [1:1000]. All antibodies were purchased from BioLegend (San Diego, CA) or BD Biosciences (San Jose, CA). Cells were surface stained for 30 minutes at room temperature. Following staining, cells were washed and fixed with BD Cytofix buffer (BD Biosciences, San Jose, CA) for 5 minutes at 4°C. Cells were washed twice and resuspended in 200 µL FACS buffer and analyzed on a BD LSR Fortessa cytometer. All data were analyzed using FlowJo software (FlowJo Inc., Ashland, OR)

### Tissue processing for immunofluorescent microscopy

Tissues to be analyzed by fluorescence microscopy were placed into 4% paraformaldehyde solution for 5 hours followed by overnight dehydration in a 30% sucrose solution at 4°C (Sigma-Aldrich, St. Louis, MO). Samples were washed with phosphate buffered saline prior to embedding within optimal cutting temperature (OCT) media (Sakura Finetek, Maumee, OH) and frozen using a dry ice slurry in 100% ethanol and stored at - 20°C. Tissues were sectioned (7 μm thick) using a cryostat (Leica CM3050 S) and affixed to positively charged Denville slides (Thomas Scientific, Swedesboro, NJ). Slides were dried at room temperature for 10 minutes preceding a 10-minute fixation in -20°C chilled 100% acetone. Slides were subsequently dried at room temperature for 10 minutes and washed twice in chilled phosphate-buffered saline before being placed in saturation buffer containing 10% normal donkey serum (Sigma-Aldrich, St. Louis, MO) and 0.1% Triton-X (Sigma-Aldrich, St. Louis, MO) in phosphate-buffered saline for blocking at room temperature for 45 minutes. Slides were incubated overnight at 4°C with a primary antibody cocktail: 1:100 mouse anti-CD45.2-AF488 (104, BioLegend, San Diego, CA), 1:200 hamster anti-CD3e-AF594 (500A2, BioLegend, San Diego, CA) and 1:200 rat anti-CD169-AF647 (3D6.112, BioLegend, San Diego, CA). The following day, slides were washed twice in chilled phosphate-buffered saline, mounted with ProLong diamond mounting media (Thermo-Fisher Scientific, Waltham, MA), covered with a coverslip, and allowed to cure overnight prior to imaging.

### Confocal microscopy and NK-cell enumeration

Confocal imaging was performed using a Laser Scanning Nikon A1RSi Inverted Confocal Microscope with NIS Elements Confocal software. Z-stacked tissue images were acquired through a 10X objective (Nikon Plan Apo λ) from which a maximum intensity projection was generated prior to cell enumeration. Using tools available in NIS Elements Analysis software and guided by CD169- and CD3-staining, we drew borders around the white pulp (CD169 boundary) and T cell zones (CD3 boundary) while visualizing only each individual channel. The channel for donor NK cell staining was hidden at this stage to prevent bias of NK cell localization on drawing of these boundaries. Thereafter, NK cells located in the previously defined T cell zones and white pulp regions were enumerated using unbiased NIS Elements-derived algorithms and plotted using GraphPad Prism (San Diego, CA). To quantify NK cells outside the white pulp, the quantity of white pulp NK cells was subtracted from the total number of NK cells detectable within a section and normalized to mm^2^. Brightness and contrast for each representative image were adjusted equally across all channels using NIS Elements Analysis.

### Statistical analysis and experimental rigor

All individual datapoints are represented in plots along with group mean, while tabular data is presented as the mean and standard deviation. Statistical differences between control and experimental groups were calculated using statistical tests indicated in each figure legend, including unpaired two-tailed Mann-Whitney (non-normally distributed data with unequal variance across two groups), unpaired two-tailed Student’s T-test (normally distributed data with equal variance across two groups), Kruskal Wallis (non-parametric comparison of two groups across time) with post-hoc Dunn’s test (quantification of *Ascl2* levels), or Dunnett’s T3 multiple comparisons Welch ANOVA (for more than two groups displaying unequal variance). A p-value of less than 0.05 was considered significant. Graphing and statistical analysis were routinely performed using GraphPad Prism (San Diego, CA). Researchers performing the tissue processing and experimental measurements were blinded to genotype and treatment conditions of the animals. This resulted in randomization of the processing order.

## Results

### Traditional CD45.1 congenic NK cells fail to thrive upon adoptive transfer

Allelic variation among inbred mouse strains at the *Ptprc* locus (encoding Ly5/CD45) has been exploited as a tool to track donor leukocytes after adoptive transfer based on the specificity of monoclonal antibodies for the *Ptprc*^*a*^ (Ly5.1/CD45.1) and *Ptprc*^*b*^ (Ly5.2/CD45.2) alleles [16]. Several groups have successfully tracked donor CD45.1+ NK cells in recipient CD45.2+ mice for extended periods of time, albeit in contexts where the donor NK cells undergo significant clonal expansion (e.g. murine cytomegalovirus infection) [11, 21] or possess homeostatic advantages over host cells (e.g. common gamma chain deficient mice) [12].

We aimed to use similar adoptive transfer model systems to study NK cell immunoregulation of T and B cells in immunocompetent mice after infection or vaccination. To our surprise, adoptive transfer of splenic NK cells from conventional B6.SJL-*Ptprc*^*a*^*Pepc*^*b*^/BoyJ (BoyJ) mice into C57BL/6J mice (**Figure 1A**) resulted in a poor or exceedingly short-lived ‘take’ (i.e. recovery from recipient tissues) of donor cells within 24 hours of cell transfer (**Figure 1B**). Based on limiting dilution studies, the expected take of donor lymphocytes (i.e. T cells) in the spleen of recipient animals is 3.1% [22]. In contrast to this expectation, donor BoyJ NK cells consistently comprised less than 0.6% of the overall pool of NK cells in recipient mice (**Figure 1C, left**), with fewer than 1,000 donor CD45.1+ NK cells recovered from host spleens (**Figure 1C, right**). Similar results were obtained with two-fold higher or lower inoculum sizes of donor NK cells (data not shown). Thus, BoyJ-derived NK cells fail to persist after transfer into immunocompetent mice, presumably due to minor histocompatibility mismatch or the effects of passenger genes located near the *Ptprc* locus [16].

**Figure 1.**
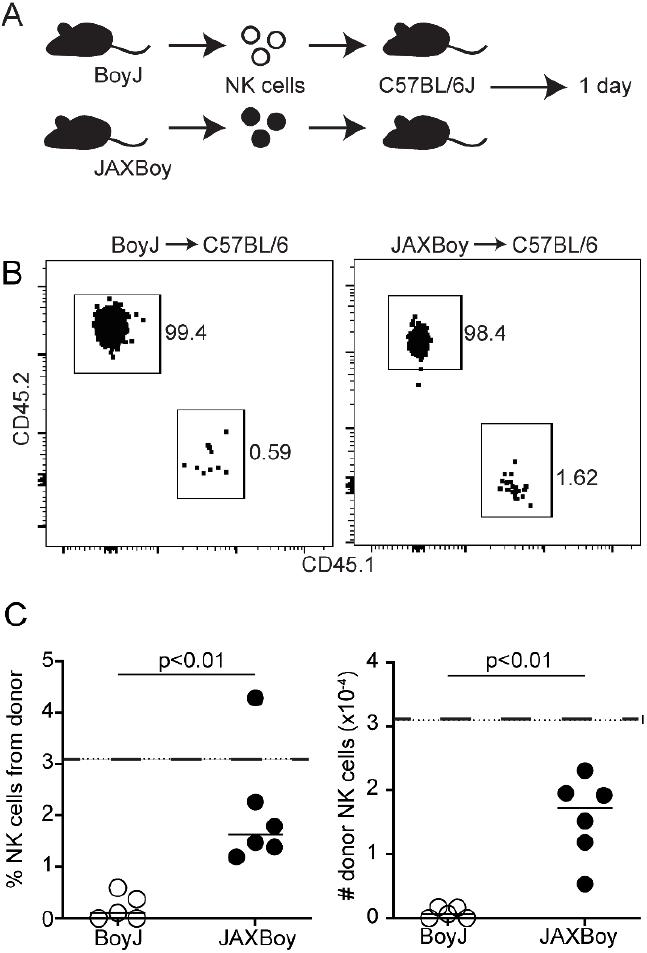
Superiority of JAXBoy over BoyJ NK cells in adoptive transfer experiments. **(A)** 1,000,000 splenic NK cells from BoyJ (CD45.1+) or JAXBoy (CD45.1+) mice were injected retro-orbitally into C57BL6/J recipients (n=5-6/group). (**B**) Representative flow plots of gated NK cells (CD4-CD8-CD19-CD49a-NK1.1+ CD49b+ NKp46+) from host (CD45.2+) or donor (CD45.1+) in recipient spleen one day after transfer. (**C**) The proportion and number of donor-derived NK cells in the spleens of each recipient mouse. Data are representative of two independent experiments. Solid lines represent mean and dotted lines represent predicted cell take (3.1%, 31,000 cells). Statistically significant differences determined by Mann-Whitney test.

### CRISPR-generated CD45.1+ coisogenic mice are a superior platform for NK transfers

The Jackson Laboratory recently developed coisogenic JAXBoy mice in which the *Ptprc*^*a*^ (Ly5.1/CD45.1) allele was introduced into C57BL/6J mice using CRISPR-Cas9 [23, 24]. This editing introduced a lysine to glutamic acid mutation at amino acid 302 (K302E; AAA->GAA) and a silent nucleotide change N303N (AAC->AAT), resulting in reactivity with anti-CD45.1 monoclonal antibodies without the confounding genetics of proximal genomic regions inherited from SJL founders in conventional BoyJ mice [16].

We transferred splenic NK cells from JAXBoy mice into C57BL/6 mice (**Figure 1A**) and analyzed recovery of donor cells from recipient mouse tissues over the ensuing five days. The average recovery of CD45.1+ JAXBoy donor NK cells in the spleen of recipient C57BL/6 mice was five times greater than that observed with BoyJ donor NK cells (**Figure 1C**). The proportion (**Figure 2A**) and number (**Figure 2B**) of CD45.1+ JAXBoy donor NK cells recovered from spleens of recipient C57BL/6 mice was relatively stable over a five-day window. Similar stability was seen in the recovery of CD45.1+ JAXBoy donor NK cells from recipient mouse lungs and liver over the same time span (**Figure 2C-F**). A reciprocal transfer of splenic CD45.2+ C57BL/6 NK into CD45.1+ JAXBoy recipient mice revealed a similarly strong take of CD45.1+ donor cells comprising an average of 1-2% of total NK and measurable numbers of donor cells in each host tissue that remained relatively stable over an experimental window of five days (**Figure 2**). In either transfer direction, donor and host splenic NK cells in the C57BL/6-JAXBoy system displayed similar phenotypic patterns of distribution across maturation states comprising immature (CD11b^-^ CD27^+^), transitional (CD11b^+^ CD27^+^), and mature (CD11b^+^ CD27^-^) subsets of cells (**Table I**) [25]. The data demonstrate that JAXBoy mice represent a superior model system for adoptive transfer studies of mouse NK cells, potentially due to the precision engineering that limits genetic effects adjacent to the *Ptprc* locus that are seen in conventional BoyJ mice [16].

**Figure 2.**
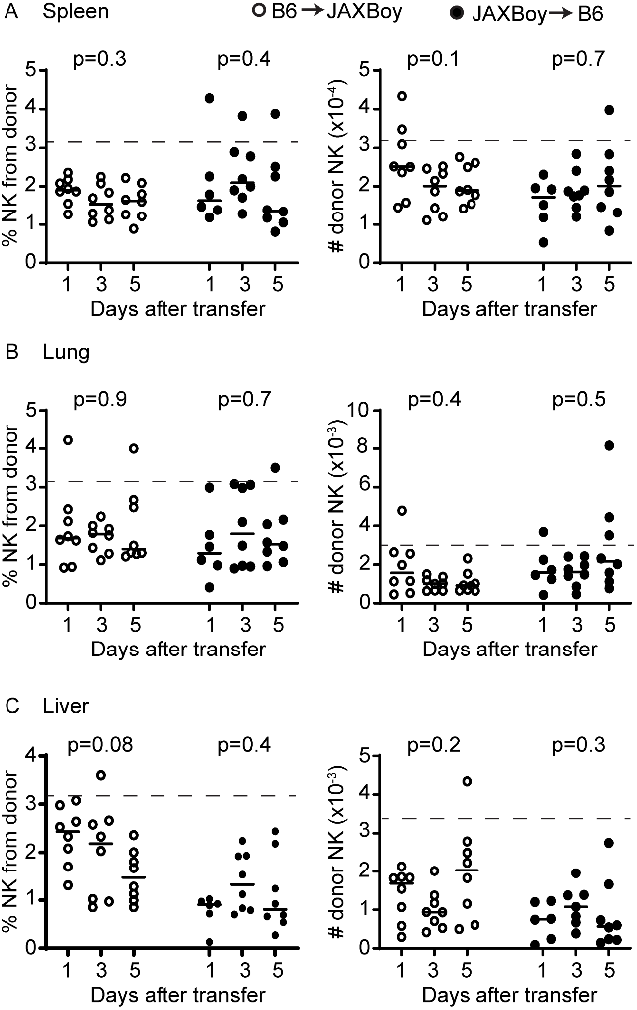
Persistence of donor NK cells in reciprocal transfer between C57BL/6 and JAXBoy mice. 1,000,000 C57BL/6 (CD45.2+, open circle) or JAXBoy (CD45.1+, closed circle) isolated splenic NK cells were transferred by retro-orbital injection into JAXBoy or B6 mice, respectively (n = 6-8/group). One, three, and five days later, recipient mouse (**A**) spleens, (**B**) lungs, and (**C**) liver were subjected to flow cytometry to determine proportion (among total CD4^-^ CD8^-^ CD19^-^ CD49a^-^ NK1.1^+^ CD49b^+^ NKp46^+^ NK cells) and number of donor-derived NK cells. Data are representative of two independent experiments. Solid lines represent mean and dotted lines represent predicted cell takes. Statistically significant differences between time points for each transfer type were determined by Kruskal-Wallis test.

**Table 1.** The percentage (mean ± standard deviation) of transferred and host NK cells (CD4^-^ CD8^-^ CD19^-^ CD49a^-^ NK1.1^+^ DX5^+^ NKp46^+^, source determined by CD45.1 (JAXBoy) or CD45.2 (C57BL/6) staining) expressing maturation markers CD11b and CD27 in the spleen 1, 3, or 5 days after transfer of 1,000,000 donor splenic NK cells. (n = 6-8/group). Data are representative of 2 independent experiments.

| JAXBoy into C57BL/6 transfer |  |  |  |  |  |  |
| --- | --- | --- | --- | --- | --- | --- |
| Days | JAXBoy donor NK cells |  |  | C57BL/6 host NK cells |  |  |
|  | CD27<sup>-</sup> CD11b<sup>+</sup> | CD27<sup>+</sup> CD11b<sup>+</sup> | CD27<sup>+</sup> CD11b<sup>-</sup> | CD27<sup>-</sup> CD11b<sup>+</sup> | CD27<sup>+</sup> CD11b<sup>+</sup> | CD27<sup>+</sup> CD11b<sup>-</sup> |
| 1 | 4.6 ± 0.02 | 21.6 ± 0.03 | 65.9 ± 0.1 | 6.8 ± 0.03 | 25.1 ± 0.1 | 60.3 ± 0.1 |
| 3 | 2.7 ± 0.02 | 20.9 ± 0.03 | 68.5 ± 0.04 | 5.6 ± 0.03 | 25.2 ± 0.1 | 59.2 ± 0.1 |
| 5 | 3.6 ± 0.01 | 14.8 ± 0.03 | 73.5 ± 0.1 | 5.0 ± 0.03 | 21.0 ± 0.1 | 66.0 ± 0.1 |
| C57BL/6 into JAXBoy transfer |  |  |  |  |  |  |
| Days | C57BL/6 donor NK cells |  |  | JAXBoy host NK cells |  |  |
|  | CD27<sup>-</sup> CD11b<sup>+</sup> | CD27<sup>+</sup> CD11b<sup>+</sup> | CD27<sup>+</sup> CD11b<sup>-</sup> | CD27<sup>-</sup> CD11b<sup>+</sup> | CD27<sup>+</sup> CD11b<sup>+</sup> | CD27<sup>+</sup> CD11b<sup>-</sup> |
| 1 | 7.0 ± 0.04 | 33.2 ± 0.1 | 53.6 ± 0.1 | 10.6 ± 0.1 | 22.4 ± 0.04 | 59.9 ± 0.1 |
| 3 | 5.4 ± 0.03 | 23.9 ± 0.1 | 59.7 ± 0.1 | 10.7 ± 0.1 | 21.2 ± 0.1 | 58.2 ± 0.1 |
| 5 | 6.1 ± 0.03 | 23.3 ± 0.1 | 62.4 ± 0.1 | 8.8 ± 0.04 | 22.1 ± 0.1 | 60.2 ± 0.1 |

### CXCR3-dependent localization of donor NK cells in splenic white pulp after infection

We recently demonstrated a transient CXCR3-dependent enrichment of NK cells in the white pulp of the spleen and lymph nodes after infection with the Armstrong strain of lymphocytic choriomeningitis virus (LCMV) [7]. This localization was required for NK cell suppression of T cell responses. To verify that donor NK cells exhibit similar CXCR3-dependent localization in recipient mice after infection [26], we transferred CD45.2+ NK cells from the spleens of C57BL/6 or *Cxcr3*-deficient (*Cxcr3* KO) mice into JAXBoy (CD45.1+) recipient mice (**Figure 3A**). Three days after cell transfer, recipient mice were intraperitoneally infected with LCMV Armstrong or left uninfected for three additional days. The recovery of CD45.2+ donor NK cells from either wild-type C57BL/6 or *Cxcr3* KO donors within uninfected and infected recipient mice was assessed via flow cytometry (**Figure 3B**). We also assessed donor (CD45.2+) NK cell positioning within the white pulp (defined by CD169+ margin surrounding a CD3+ T cell zone) using confocal microscopy at day three of infection (**Figure 3C**). Infection triggered a statistically significant increase in the number of wild-type donor CD45.2+ (C57BL/6) NK cells in both the white pulp (**Figure 3D, left**) and T-cell zones (**Figure 3D, right**) of JAXBoy recipients relative to uninfected control hosts. No statistically significant enrichment of *Cxcr3* KO donor CD45.2+ NK cells was observed in either the white pulp (**Figure 3E, left**) or T-cell zones (**Figure 3E, right**) of recipient JAXBoy mice after virus infection. Given the vanishingly small number of NK cells present in lymph nodes in prior studies [7], we focused the present work exclusively on donor cell localization in the spleen. By subtracting white pulp localized NK cells from the total, we estimated the number of NK cells in other regions of the spleen (**Figure 3F**). This analysis revealed similar numbers of WT NK cells in non-white pulp regions before and after infection, while Cxcr3 KO NK cells were enriched in non-white pulp regions only after infection. These results confirm prior studies demonstrating CXCR3-dependent transient enrichment of NK cells in the white pulp regions of the spleen after infection [7], yet suggest that CXCR3 is not required for infection-induced accumulation of NK cells in the spleen. Moreover, these data highlight the utility of this NK-cell transfer model to aid the discovery of additional mediators governing the critical positioning of NK cells.

**Figure 3.**
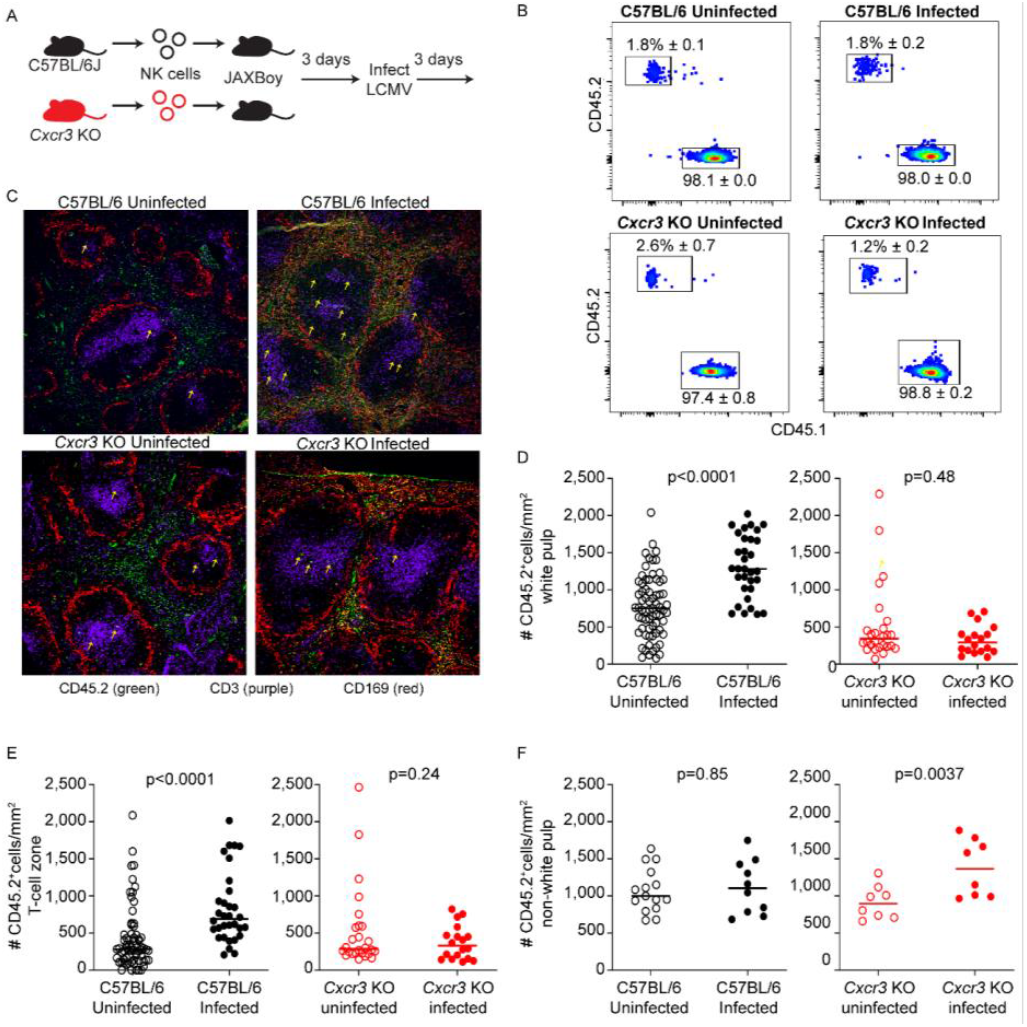
CXCR3 is required for donor NK cells to localize in spleen white pulp and T-cell zones after infection. **(A)** 1,000,000 CD45.2+ C57BL/6 or *Cxcr3* KO NK cells were retro-orbitally injected into JAXBoy (CD45.1+) recipient mice (n = 3/group). Three days after transfer, recipient mice were infected with LCMV or left uninfected. (**B**) Three days later, take of donor cells (CD45.2+) was confirmed via flow cytometry in recipient mice (CD45.1+). (**C**) CD45.2+ (green) donor NK cell positioning within spleen relative to CD169+ marginal zone macrophages (Red) and CD3+ T cells (Purple) assessed by confocal microscopy for both C57BL/6 and *Cxcr3* KO donor NK cells. Arrows demarcate representative donor NK cell positioning within these sections. The relative positioning of C57BL/6 or *Cxcr3* KO donor NK cells in the (**D**) CD169+ bound white pulp or (**E**) CD3+ T-cell zones was quantified in 10-12 follicles per recipient mouse (horizontal line represents mean). Open symbols reflect uninfected mice, and closed symbols reflect mice infected with LCMV. (**F**) The number of C57BL/6 or *Cxcr3* KO donor NK cells outside the white pulp was calculated for each image by subtracting the number of NK cells quantified within the CD169-defined white pulp from the total number of NK cells in the image. Data are representative of 2 independent staining experiments. Statistical significance differences in donor NK cell densities in various spleen regions between uninfected and infected host mice were determined by Mann Whitney analysis.

### Role of ASCL2 in NK cell migration to spleen white pulp during infection

Achaete-Scute Complex Homolog 2 (ASCL2) is a transcription factor implicated in the differentiation of follicular helper T cells and their localization within lymphoid tissues [27]. ASCL2 transcripts are measurably expressed in mouse [28] and human [29] NK cells, but the function of this gene in innate lymphocytes remains unexplored. We hypothesize that NK cells may require ASCL2 for precise localization to subdomains of lymphoid tissues during infection (i.e. infiltration of white pulp). We generated mice in which NK cells lack *Ascl2* by crossing *Ncr1*-*Cre* [18] animals with mice in which the second exon of *Ascl2* is flanked by LoxP sites [19]. The resulting *Cre*+ conditional *Ascl2*-deficient mice (*Ascl2* cKO) were viable, fertile, and contained statistically similar proportions and numbers of NK cells in their spleens in comparison to control *Ascl2-floxed Cre*-negative (*Ascl2* WT) littermates (**Figure 4A-B**). Of note, the MFI of NKp46 expression is reduced in *Ascl2* cNK cells due to hemizygous expression of NKp46 in *Ncr1-iCre+* animals, as previously described [18].

**Figure 4.**
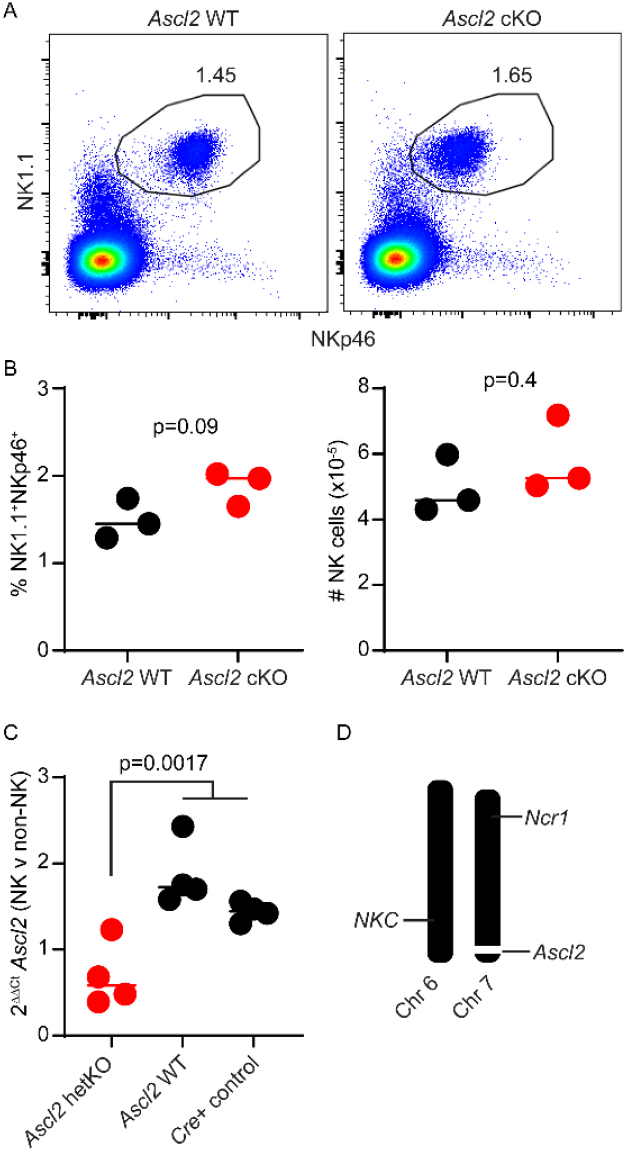
*Ascl2* is dispensable for the development of mouse NK cells. **(A)** Spleens from *Ascl2* WT (*Cre*-negative *Ascl2*^*fl/fl*^,) and *Ascl2* conditional knockout (cKO, *Cre*+ *Ascl2*^*fl/fl*^) mice (n=3/group) were collected and stained to quantify NK cells by flow cytometry. (**B**) The percentage and number of NK cells (CD3-NK1.1+NKp46+) in the spleen is quantified and plotted for *Ascl2* WT (black) and *Ascl2* cKO (red). The horizontal line represents median. Statistically significant differences were determined by unpaired Student’s t-test. Data are representative of 2 independent staining experiments. (**C**) NK cells were isolated by positive selection and corresponding non-NK cells were collected from the spleens of *Ncr1-Cre*+ *Ascl2*^*fl/WT*^ (*Ascl2* hetKO, red), *Ncr1-Cre*(-) *Ascl2*^*fl/WT*^ (*Ascl2* WT, black), and *Ncr1-Cre*+ control mice (black). Quantitative PCR amplification of *Ascl2* and *Gapdh, Ascl2* Ct normalized to *Gapdh* (ΔCt), NK cell ΔCt was normalized to non-NK ΔCt (ΔΔCt), and fold-change transposition (2^-ΔΔCt^) were performed. Significant differences between genotypes assessed by Kruskal-Wallis with a post-hoc uncorrected Dunn’s test. (**D**) Visual representation of SNP genotyping analysis (Supplemental Table I) for key chromosomes in *Ascl2*-floxed mouse colony, where black is homozygous for C57BL/6 SNP and white bars equate with homozygous for non-C57BL/6 SNP. Important NK cell receptor gene complex (NKC) on chromosome 6 and *Ncr1* (NKp46) region on chromosome 7 are homozygous C57BL/6, while locus around floxed *Ascl2* gene on chromosome 7 is not C57BL/6-derived as expected.

Although detection of ASCL2 protein proved challenging, we confirmed by real-time quantitative PCR that *Ascl2* mRNA expression was selectively reduced in NK cells but not in other immune cells of *Ascl2* cKO mice relative to control mice (**Figure 4C**). Since *Ascl2* genetic targeting was initially performed in 129/Ola mouse-derived embryonic stem cells, we performed genome-wide SNP genotyping on our colony. After more than 8 backcrosses to C57BL/6, the *Ascl2*-floxed mice were >96% C57BL/6 (**Supplemental Table 1**), expressed key C57BL/6-associated NK cell receptors NK1.1 and Ly49H (not shown), and displayed homozygous C57BL/6 congenic genotype across chromosomes 6 (site of NK cell receptor gene complex) and *Ncr1-*end of chromosome 7 while exhibiting non-C57BL/6 DNA proximal to the floxed *Ascl2* alleles (**Figure 4D** and **Supplemental Table 1)**.

We intravenously transferred splenic CD45.2+ NK cells from either *Ascl2* cKO or *Ascl2* WT mice into JAXBoy mice three days before infection with LCMV Armstrong (**Figure 5A**). The recovery of donor CD45.2+ *Ascl2* WT or *Ascl2* cKO NK cells in the spleen of recipient CD45.1+ JAXBoy mice was assessed by flow cytometry (**Figure 5B**). We analyzed the positioning of donor NK cells in recipient mouse spleens on day three of infection and in uninfected hosts (**Figure 5C**). Infection triggered a statistically significant increase in the number of *Ascl2* WT donor NK cells present within the white pulp or T-cell zones (**Figure 5D-E**) compared to uninfected mice. In contrast, *Ascl2* cKO NK cells were not enriched in the white pulp or T-cell zones after infection (**Figure 5D-E**). In contrast to loss of Cxcr3 in **Figure 3**, the accumulation of NK cells in total spleen after infection was only observed for *Ascl2* WT but not *Ascl2* cKO NK cells (**Figure 5F**). These data indicate that *Ascl2* is important for infection-induced accumulation of NK cells in the splenic white pulp potentially by regulating their recruitment to the spleen itself.

**Figure 5.**
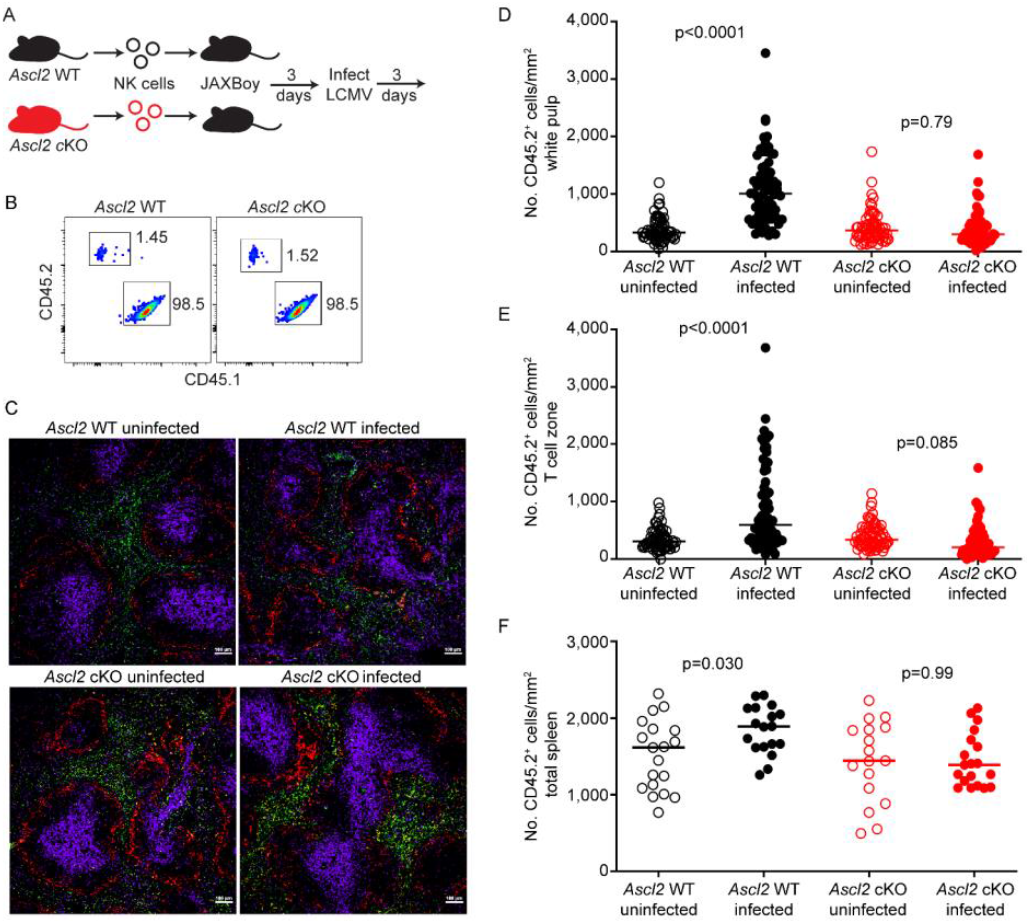
*Ascl2* is required for NK cell positioning in splenic white pulp after infection. (**A**) *Ascl2* WT or *Ascl2* cKO NK cells (CD45.2+) NK cells were adoptively transferred into JAXBoy (CD45.1+) mice (n = 2/group) 3 days prior to infection with LCMV or being left uninfected for an additional 3 days. (**B**) The take of CD45.2+ donor cells was confirmed via flow cytometry in uninfected CD45.1+ recipient mouse spleens. (**C**) CD45.2+ (green) donor NK cells were quantified based on positioning within spleen using microscopy relative to CD169+ marginal zone macrophages (Red) and CD3+ T cells (Purple). Representative images at 10x magnification are shown. (**D-F**) The densities of CD45.2+ *Ascl2* WT (black circles) or *Ascl2* cKO (red circles) donor NK cells in specific splenic regions of uninfected (open circles) or infected (closed circles) host mice were quantified in 10-12 follicles per recipient mouse (horizontal line represents mean). Regions include (**D**) CD169+ bound white pulp, (**E**) CD3+ T-cell zones, and (**F**) total spleen. Data are representative of 2 independent staining experiments. Statistically significant differences between uninfected and infected donor NK cell densities in various spleen regions were determined by post-hoc Dunnett’s T3 multiple comparisons testing with Welch ANOVA (latter p values are shown).

## DISCUSSION

These results demonstrate the improved utility of CRISPR-generated JAXBoy mice in comparison to conventional BoyJ mice for adoptive transfer and tracking of donor NK cells in immunocompetent recipient animals. This distinction is likely due to the limited number of genetic variants (two nucleotides) introduced into JAXBoy mice to confer CD45.1 reactivity on the C57BL/6 background. In contrast, prior work revealed large and variable co-inheritance of non-C57BL/6 alleles (SJL derived) near the *Ptprc* locus in conventional BoyJ mice [16]. These include genes that affect survival (*Bcl2*), migration (*Cxcr4, Sell*), antiviral interferon responses (*Rnasel*), and immune function (*Pdcd1, Il10*), any of which are likely important in the survival and positioning of NK cells in cell transfer model systems. The JAXBoy mouse likely holds similar advantages over conventional BoyJ mice in adoptive transfer experiments with other types of CD45-expressing leukocytes. The difference seen in donor cell uptake in our experiments also raises concerns for the use of conventional CD45.1+ congenic transfer mice in other experimental systems, including those where donor T cell or NK cell proliferation may mask these drawbacks.

We present a new cell transfer model applicable to the investigation of intrasplenic migration of NK cells. We use this model to confirm the necessity of CXCR3 for positioning of NK cells in the white pulp near T and B cells after virus infection. Yet, our data additionally indicates that CXCR3 is dispensable for the general accumulation of NK cells in the spleen during infection. The mispositioning of CXCR3-deficient NK cells in the red pulp likely explains the inability of these CXCR3-less cells to regulate T cell responses observed in our prior publication [30]. Furthermore, we develop an original strain of mice with conditional deletion of Ascl2 in NK cells and employ this new transfer model to demonstrate a role for ASCL2 in enrichment of NK cells in the spleen and white pulp after virus infection. ASCL2 controls the expression of numerous target genes involved in follicular T cell [27, 31] and germinal center B cell [32] biology, including cell migration receptors CXCR5 and CXCR4. ASCL2 also intersects with other transcriptional networks implicated in various facets of NK cell biology, including ETS1 [32] and BLIMP1 [31]. Thus, the discovery of a role for ASCL2 in NK cell positioning in the spleen is the first step in a new area of research into the role of this transcriptional regulator in NK cell biology. This and future studies are likely to reveal ASCL2-regulated pathways vital for immunoregulatory functions of NK cells that can be targeted to circumvent NK cell suppression of vaccine response or to bolster NK-cell based therapies targeting pathogenic T and B cells in autoimmune disease.

## Acknowledgements

We thank Drs. Hans Clevers and James Bridges for providing initial breeder pairs of Ascl2-floxed mice. All flow cytometric data were acquired using equipment maintained by the Research Flow Cytometry Facility in the Division of Rheumatology at Cincinnati Children’s Hospital Medical Center. The confocal microscopy data was made possible, in part, using the Cincinnati Children’s Bio-Imaging and Analysis Facility [RRID: SCR_022628].

## Author contributions

Conceptualization and design of study (LMC, SMW), performance of experiments and data acquisition (LMC, AC, HAF, HS, AA, JAT, LBC, SNB, AY, MW), data analysis (LMC, SNW), drafting of the manuscript (LMC, SNW), and critical editing of the manuscript (LMC, AC, HAF, HS, AA, JAT, LBC, SNB, AY, MW).

